# A comparative analysis among ADAR mutant mice reveals site-specific regulation of RNA editing

**DOI:** 10.1101/822916

**Authors:** Pedro Henrique Costa Cruz, Yuki Kato, Taisuke Nakahama, Toshiharu Shibuya, Yukio Kawahara

## Abstract

Adenosine-to-inosine RNA editing is an essential posttranscriptional modification catalyzed by adenosine deaminase acting on RNA (ADAR)1 and ADAR2 in mammals. For numerous sites in coding sequences (CDS) and microRNAs (miRNAs), editing is highly conserved and has significant biological consequences, for example, by altering amino acid residues and target recognition. However, technical limitations have prevented a comprehensive and quantitative study to determine how specific ADARs contribute to each site. Here, we developed a simple method in which each RNA region with an editing site was amplified separately and combined for deep sequencing. Using this method, we compared the editing ratios of all sites that were either definitely or possibly conserved in CDS and miRNAs in the cerebral cortex and spleen of wild-type mice, *Adar1^E861A/E861A^Ifih^−/−^* mice expressing inactive ADAR1 (Adar1 KI) and *Adar2^−/−^Gria2^R/R^* (Adar2 KO) mice. We found that the editing ratio was frequently upregulated in either Adar mutant mouse strain. In contrast, we found that the presence of both ADAR1 and ADAR2 was required for the efficient editing of specific sites. In addition, some sites, such as miR-3099-3p, showed no preference for either ADAR. We further created double mutant Adar1 KI Adar2 KO mice and observed viable and fertile animals with complete absence of editing, suggesting that ADAR1 and ADAR2 are the sole enzymes responsible for all editing sites *in vivo*. Collectively, these findings indicate that editing is regulated in a site-specific manner by the different interplay between ADAR1 and ADAR2.

## INTRODUCTION

RNAs are known to be subjected to multiple post-transcriptional modifications that affect each’s fate. One such RNA modification is adenosine-to-inosine (A-to-I) RNA editing, which is widely conserved among many organisms, from nematodes to humans (Hundley and Bass 2010; Savva et al. 2012; Nishikura 2016). Although RNA editing sites were previously found by chance, the number of sites identified has dramatically increased after the introduction of deep sequencing technology (Li et al. 2009; Ramaswami et al. 2013; Sakurai et al. 2014), with an estimate of more than 100 million RNA editing sites present in human transcripts (Bazak et al. 2014; Picardi et al. 2015; Tan et al. 2017). Most of these sites are located in repetitive elements (REs) in non-coding regions, given that REs occasionally form double-stranded RNA (dsRNA) structures, which are required for adenosine-to-inosine RNA editing. As a consequence, RNA editing in REs alters dsRNA structure, which is indispensable in escaping recognition as non-self by the host immune system (Mannion et al. 2014; Liddicoat et al. 2015; Pestal et al. 2015). Although this role is conserved among mammals, RNA editing in REs occurs frequently in primates due to the abundance of REs, especially primate-specific Alu repeats (Neeman et al. 2005); it therefore follows that the total number of editing sites is much lower in rodents (Danecek et al. 2012; Li and Church 2013). In contrast, although their frequency is quite rare, RNA editing sites in protein-coding sequences (CDS) and microRNAs (miRNAs) are relatively conserved among mammals (Li et al. 2009; Li and Church 2013; Pinto et al. 2014; Nishikura 2016; Jinnah and Ulbricht 2019). Given that inosine is recognized as if it were guanosine by the translational machinery, RNA editing in CDS leads to re-coding events that can potentially affect the functions of the corresponding proteins (Pullirsch and Jantsch 2010). Indeed, mutant mice expressing either an edited or unedited protein alone exhibit abnormal phenotypes, such as being more prone to seizures, hyperactivity, depression and dysregulated vascular contractions (Higuchi et al. 2000; Kawahara et al. 2008a; Mombereau et al. 2010; Miyake et al. 2016; Jain et al. 2018). In addition, dysregulated RNA editing in CDS is linked to human diseases such as cancer and amyotrophic lateral sclerosis (Kawahara et al. 2004; Slotkin and Nishikura 2013; Peng et al. 2018). Furthermore, editing in miRNAs affects their expression and target recognition (Yang et al. 2005; Kawahara et al. 2007a; Kawahara et al. 2007b; Kawahara et al. 2008b). Therefore, it is crucial to know the enzyme(s) responsible for each conserved site in order to understand the mechanisms underlying the tight regulation of RNA editing *in vivo*.

Adenosine deaminases acting on RNAs (ADARs) are the enzymes responsible for adenosine-to-inosine RNA editing, which requires a dsRNA structure for target recognition. In mammals, three ADARs have been identified (Hundley and Bass 2010; Nishikura 2016): while brain-specific ADAR3 appears enzymatically inactive, ADAR1 and ADAR2 are active enzymes that are ubiquitously expressed, although their expression level varies in a tissue-specific manner (George et al. 2005; Picardi et al. 2015; Huntley et al. 2016; Heraud-Farlow et al. 2017; Tan et al. 2017; Nakahama et al. 2018). For instance, ADAR2 is mainly localized in the nucleus, and highly expressed in the brain and aorta (Huntley et al. 2016; Tan et al. 2017; Jain et al. 2018; Nakahama et al. 2018). In contrast, ADAR1 has two isoforms: a short p110 isoform that is mainly localized in the nucleus and is highly expressed in the brain, and a long p150 isoform that is mainly localized in the cytoplasm and is highly expressed in the thymus and spleen, where ADAR2 is expressed at low level (Huntley et al. 2016; Nakahama et al. 2018). In addition to the different expression patterns of ADARs, it is known that ADAR1 and ADAR2 regulate RNA editing in a competitive manner in some cases, which has made it difficult to determine the contribution of each ADAR to the RNA editing of each conserved site *in vivo* (Kawahara et al. 2007b; Riedmann et al. 2008; Wahlstedt et al. 2009; Vesely et al. 2014; Picardi et al. 2015; Huntley et al. 2016; Tan et al. 2017). One simple solution to this problem may be by comparing the editing ratio of each site among wild-type (WT), Adar1-deficient (*Adar1^−/−^*) and Adar2-deficient (*Adar2^−/−^*) mice. However, *Adar1^−/−^* and *Adar2^−/−^* mice exhibit embryonic and early postnatal lethality, respectively (Higuchi et al. 2000; Hartner et al. 2004; Wang et al. 2004). Fortunately, the lethality of *Adar2^−/−^* mice can be rescued by the single substitution of adenine to guanine in an editing site in the coding region of *Gria2* at the genomic DNA level. This leads to a change in one amino acid residue, from glutamine (Q) to arginine (R), which allows *Adar2^−/−^Gria2^R/R^* mice (which we termed Adar2 KO mice) to survive until adulthood (Higuchi et al. 1993). In contrast, although the embryonic lethality of *Adar1^−/−^* mice can be rescued by concurrent knockout of the *Ifih1* gene encoding MDA5, a cytoplasmic sensor for dsRNAs, *Adar1^−/−^Ifih^−/−^* mice still develop postnatal lethality for unknown reasons (Pestal et al. 2015). Therefore, several studies have tried to determine the ADAR responsible for each site by comparing primary neuronal cultures prepared from *Adar1^−/−^* mice and the brains of *Adar2^−/−^* mice (Riedmann et al. 2008), embryos collected from different *Adar* mutant mice (Vesely et al. 2012; Tan et al. 2017), or HeLa cells in which either ADAR1 or ADAR2 was knocked down (Nishimoto et al. 2008). However, in addition to the difficulties in comparing the different conditions, editing activity is relatively low in embryos and cultured cells. Of note, an editing-inactive E861A point mutation (*Adar1^E861A/E861A^* mice) in mutant mice is embryonically lethal, whereas it was recently reported that *Adar1^E861A/E861A^Ifih^−/−^* mice (which we termed Adar1 KI mice) survive with a normal life-span (Liddicoat et al. 2015; Heraud-Farlow et al. 2017). Therefore, it is now possible to determine the contribution of each ADAR to the editing of conserved sites by comparing the editing ratio between WT, Adar1 KI and Adar2 KO mice. However, although it is advantageous for the comprehensive identification of editing sites, total RNA-sequencing (RNA-seq) analysis is a poor method for accurate quantification of the editing ratio due to its limited sequencing depth (Tan et al. 2017). To compensate for this disadvantage, a microfluidic multiplex PCR and deep sequencing (mmPCR-seq) method was developed, in which PCR is performed with multiplex primers targeting each editing site followed by deep sequencing (Zhang et al. 2013). This method can quantify the editing ratio at many sites more accurately, although the amplification efficiency of each target is affected by the expression level of each RNA and primer design, which implies a lack of guarantee in obtaining the editing ratio of the sites targeted. In this regard, although the global dynamics of RNA editing (mostly in REs) were recently revealed by comparing the editing ratios between WT, Adar1 KI and Adar2 KO mice using the mmPCR-seq method for a limited number of samples (1 WT mouse vs. 2 Adar1 KI mice), none of the sites in CDS were identified as an ADAR1 site in the brain, for example (Tan et al. 2017). In addition, this analysis did not examine any editing sites in miRNAs. Therefore, it remains challenging to develop a method that comprehensively and accurately quantifies the editing ratio at conserved sites, which is important for understanding the contribution of each ADAR.

In this study, we developed a method in which reverse-transcription (RT)-PCR was performed for each RNA editing site followed by an adjustment of the amplicon length through a second round of PCR. After gel-purification of each PCR product, similar amounts were combined for deep sequencing. Although this method is laborious, it yields editing ratios for all sites examined with a high degree of accuracy. We applied this method to all RNA editing sites in CDS and miRNAs that are definitely or possibly conserved between humans and mice, and compared editing ratios in the cerebral cortex and spleen between WT, Adar1 KI and Adar2 KO mice. Through this analysis, we found that a considerable number of sites revealed a higher editing ratio in either Adar1 KI or Adar2 KO mice than that in WT mice, which suggests that ADARs competed with each other in these cases. In contrast, we identified some sites, such as the serotonin (5-HT) 5-HT2CR receptor (5-HT_2C_R) B site, that required a coordinated interplay between ADAR1 and ADAR2 for efficient editing. In addition, editing was preserved in both ADAR1 KI and ADAR2 KO mice at several other sites, such as miR-3099-3p, which suggests a lack of preference for either ADAR. We also established Adar1 KI Adar2 KO mice for the first time and found that RNA editing was completely absent, which suggests that ADAR1 and ADAR2 are the sole editing enzymes *in vivo*. These findings indicate that RNA editing is regulated in a site-specific manner through the different interplay between ADAR1 and ADAR2, and that the relative dosage of each ADAR is a factor that underlies the tight regulation of site-specific RNA editing. In addition, our comprehensive and quantitative data will be a valuable resource for identifying the contribution of each ADAR to all conserved sites.

## RESULTS AND DISCUSSION

### Editing ratios for CDS and miRNA sites in the cerebral cortex are higher than for the spleen with a few exceptions

To comprehensively and accurately quantify editing ratios in conserved editing sites in humans and mice, we developed a simple method in which each target site was separately amplified by RT-PCR (Fig. 1A). After adjusting the length of PCR products within 190–200 bp and adding adaptor and barcode sequences with a second round of PCR, each PCR product was gel-purified. After measuring the concentration of each sample, 30–200 PCR products were combined using approximately equal amounts and then subjected to deep sequencing (Fig. 1A). This method was laborious, but has the advantage of quantifying the editing ratio at all sites to be investigated with high precision, regardless of the expression level of each target RNA. To list the conserved editing sites in CDS and miRNAs, we initially selected A-to-I RNA editing sites reported to be conserved in humans and mice (Kawahara et al. 2008b; Chiang et al. 2010; Maas et al. 2011; Alon et al. 2012; Danecek et al. 2012; Ekdahl et al. 2012; Gu et al. 2012; Daniel et al. 2014; Pinto et al. 2014; Ramaswami and Li 2014; Vesely et al. 2014; Nishikura 2016; Terajima et al. 2016). Next, by Sanger sequencing we preliminarily examined RNA editing in the cerebral cortex and spleen for sites reported to be possibly edited in mice and that may be conserved in humans (HIST2H2AB L/L, HIST2H2AC N/S, ZNF397 N/D, miR-542-3p, miR-574-5p and miR-708-3p) (Cattenoz et al. 2013; Vesely et al. 2014; Hosaka et al. 2018). We observed possible RNA editing only in HIST2H2AB L/L and HIST2H2AC N/S sites, which were therefore included in the list. We further preliminarily examined RNA editing for sites reported to be present only in humans but that are possibly conserved in mice (AR T/A, RHOQ N/S, NCSTN S/G, TNRC18 E/G, XKR6 R/G, BEST1 I/V, GIPC1 T/A and GIPC1 P/P sites, and miR-200b and miR-455) (Martinez et al. 2008; Han et al. 2014; Sakurai et al. 2014; Nishikura 2016; Wang et al. 2017). We subsequently detected possible RNA editing in only miR-200b and miR-455, which were included in the list. Finally, the NEIL1 K/K site was reported to be present in only human cancer cells (Anadón et al. 2015). However, this site is adjacent to a conserved K/R site, and therefore was included as a possible conserved site in the list. Consequently, we listed 69 sites in the CDS of 39 genes and 26 sites in 21 miRNAs as sites that were definitely or possibly conserved, and these were used for subsequent analysis. In addition, some representative REs in the intron and 3’ untranslated region (3’UTR) (Nakahama et al. 2018), and in intronic self-editing sites in the *Adar2* gene (Rueter et al. 1999) were included as references. We then selected the cerebral cortex and spleen as representative tissues for this analysis because, as previously reported (Nakahama et al. 2018), ADAR1 p110 and ADAR2 are highly expressed in the cerebral cortex while cytoplasmic ADAR1 p150 is undetectable, and ADAR1 p150 is highly expressed in the spleen while ADAR2 is expressed at very low levels (Fig. 1B). PCR products containing the editing sites were amplified from these two tissues isolated from male WT, Adar1 KI and Adar2 KO mice at 8 weeks of age (n = 3 mice for each group). We found that the inactivation of ADAR1 or deletion of ADAR2 did not induce a compensatory upregulation of the remaining ADAR (Fig. 1B). Compensatory upregulation of *Adar2* mRNA was also not reported in the brain and spleen of Adar1 KI mice (Heraud-Farlow et al. 2017). We successfully obtained editing ratios for all sites examined from the cerebral cortex (Supplemental Table S1). In contrast, we did not determine editing ratios at 18 sites in 11 genes in the spleen due to a lack of amplification of PCR products since most of these genes are expressed in a tissue-specific manner, such as the brain-specific *Htr2c* gene (Kawahara et al. 2007b; Kawahara et al. 2008a). First, to validate our method, we focused on two representative editing sites, i.e. AZIN1 S/G, a known ADAR1 site (Chen et al. 2013), and Kv1.1 I/V, a known ADAR2 site (Bhalla et al. 2004). We did not detect editing at the AZIN1 S/G site in the spleen of Adar1 KI mice, and no significant difference was observed between WT and Adar2 KO mice (Fig. 1C). In contrast, we found no significant difference in the editing ratio at the Kv1.1 I/V site in the cerebral cortex between WT and Adar1 KI mice, although the editing ratio was dramatically reduced to 1% in Adar2 KO mice (Fig. 1C). These results clearly suggest that ADAR1 and ADAR2 are the sole enzymes responsible for AZIN1 S/G and Kv1.1 I/V sites, respectively, and support our methodology. We further validated our methodology by randomly selecting 104 PCR products containing various editing sites from all mice examined and subjected these to resequencing. This analysis underscored the high reproducibility of technical replicates (Supplemental Table S2), which suggested that the calculation of the editing ratio was not affected by the combination of PCR products for sequencing and the read number, at least if the minimum threshold for the number of total reads was set to 1,000 as described in Material and Methods. In addition, we then prepared PCR products again for 40 randomly selected targets from the total RNA used in the first analysis, which were then subjected to sequencing. This analysis also revealed a high correlation between two independent analyses of the same RNAs (Supplemental Table S3), which further strengthened the validity of our methodology. We then compared the editing ratios of all sites examined in the cerebral cortex and spleen in WT mice. We found that editing ratios in most CDS and miRNA sites in the cerebral cortex were higher than in the spleen (Fig. 1D). This is in line with these sites being predominantly edited in the nucleus. More specifically, editing in the CDS requires an editing (or exon) complementary sequence (ECS), which is usually located in an adjacent intron (Gerber and Keller 2001); therefore, these sites can be edited only in the nucleus. In contrast, the editing ratio of REs in the 3’UTR was higher in the spleen, which is likely attributed to the contribution of ADAR1 p150 in the cytoplasm. Of note, some sites in the CDS were edited at a higher level in the spleen. In particular, AZIN1 E/G and S/G sites, which are ADAR1 sites, were edited by 10% and 19%, respectively, in the spleen, whereas these sites were edited by less than 1% in the cerebral cortex of WT mice where ADAR1 p110 is highly expressed (Fig. 1C-D). These results suggest that the ECS of AZIN1 E/G and S/G sites might be located in the same exon but not in the adjacent intron. Indeed, although it was previously reported that the estimated dsRNA structure is formed with a part of the downstream intron (Chen et al. 2013), the formation of a dsRNA structure only within the exon is also possible (Supplemental Fig. S1A). Furthermore, we compared the editing ratios of AZIN1 E/G and S/G sites between mature mRNA and precursor mRNA (pre-mRNA) in the spleen, which showed that only mature mRNA was edited at both sites (Supplemental Fig. S1B). Therefore, considering that the ECSs of the Kv1.1 I/V and the GABRA3 I/M sites are located within the same exon (Bhalla et al. 2004; Rula et al. 2008), AZIN1 E/G and S/G sites may be an additional case (and the first case as ADAR1 sites) in which a dsRNA structure forms within a single exon.

**FIGURE 1.**
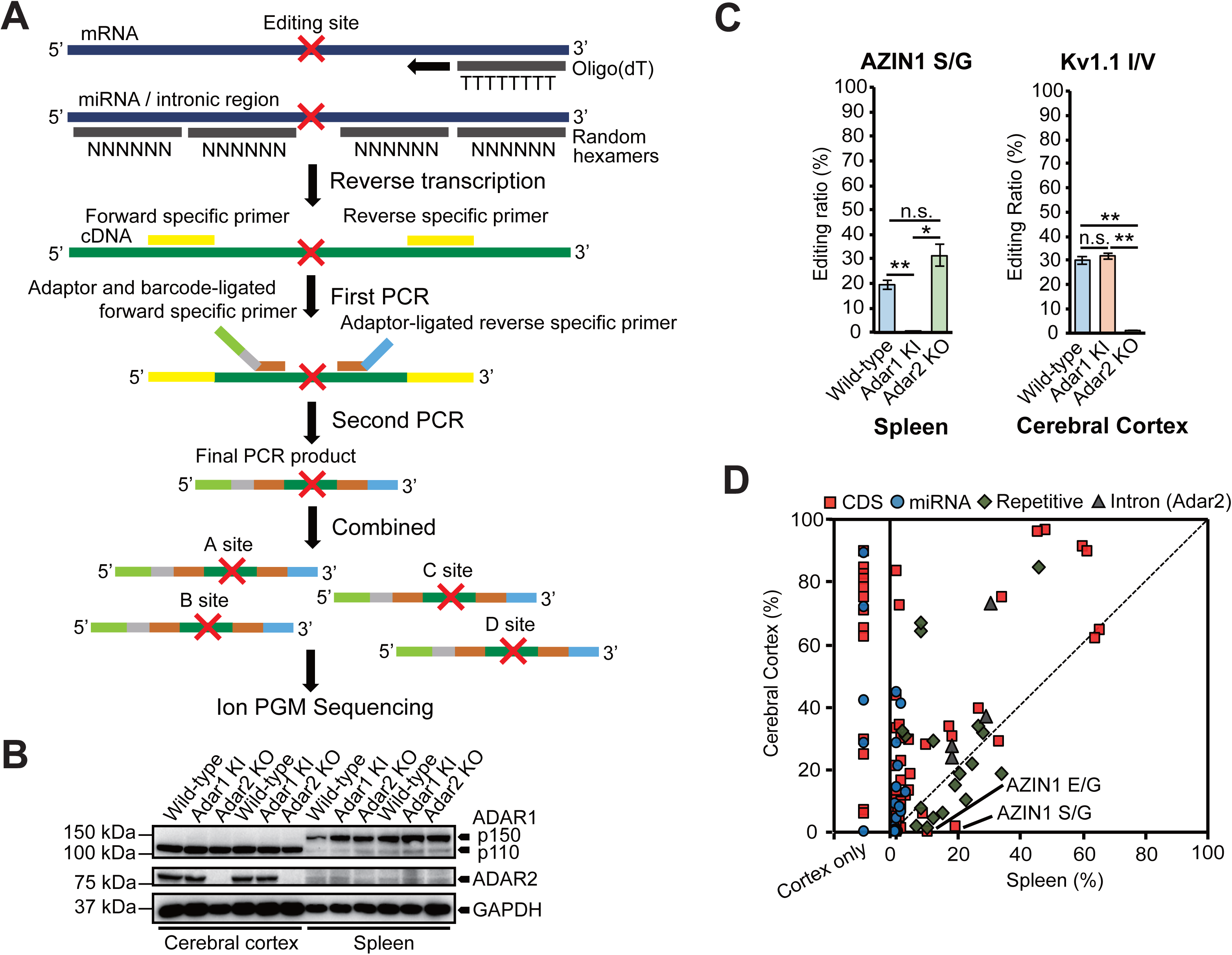
Comparison of RNA editing ratios between the cerebral cortex and spleen of WT mice. (*A*) The protocol to create Ion amplicon libraries for the evaluation of RNA editing ratios at multiple sites. After reverse-transcription using oligo(dT) primers or random hexamers, the first PCR was performed using cDNA (in green) that included an RNA editing site (shown as a red cross) and first primers specific for each editing site (in yellow). Then, a second round of PCR was performed using an aliquot of the first PCR product as a template, with each second forward primer specific to the editing site and containing an A adaptor (in light green), an Ion Xpress Barcode™ (in gray) and editing site-specific sequences (in brown), and a reverse primer that contained a trP1 adaptor (in light blue); editing site-specific sequences (in brown) were also included. All second PCR products were designed to be 190 to 200 bp in length. After 50–300 PCR products were combined, the samples were sequenced using an Ion Torrent™ Personal Genome Machine™ (Ion PGM) system. (*B*) Immunoblot analysis of adenosine deaminase acting on RNA (ADAR)1 p110, ADAR1 p150 and ADAR2 expression in cerebral cortexes and spleens isolated from wild-type (WT), *Adar1^E861A/E861A^Ifih^−/−^* mice (Adar1 KI) and *Adar2^−/−^Gria2^R/R^* (Adar2 KO) mice (n=2 mice for each group). The expression of GAPDH is shown as a reference. (*C*) Validation of the methodology by referring to the editing ratios of known ADAR1 (AZIN1 serine/glycine [S/G]) and ADAR2 sites (Kv1.1 isoleucine/valine [I/V]). Editing ratios at each site in each indicated tissue isolated from WT, Adar1 KI and Adar KO mice are displayed as the mean ± SEM (n=3 mice for each group; Student’s *t*-test, \**p* < 0.05, \*\**p* < 0.01, n.s., not significant). (*D*) Editing ratios of all sites examined were compared between cerebral cortexes and spleens isolated from WT mice. Values are displayed as the mean of values from three mice. The red squares, blue circles, green diamonds and grey triangle dots represent editing sites in coding sequences (CDS), microRNAs (miRNAs), repetitive elements and introns, respectively. Editing ratios for sites that could only be amplified from the cerebral cortex are separately displayed in the “Cortex only” fraction.

### Upregulation of editing is frequently observed in non-dominant ADAR mutant mice

Next, we examined the editing retention rate in Adar1 KI and Adar2 KO mice. In most sites, the editing ratio was greatly reduced in either Adar1 KI or Adar2 KO mice, which suggested that one Adar dominantly contributed to the editing of these sites (Fig. 2A-B). In contrast, unexpectedly, we found that the editing ratio was frequently increased in non-dominant Adar mutant mice, especially in the spleen of Adar2 KO mice. This was not caused by a difference in the presence of inactive ADAR1 in Adar1 KI mice and the total absence of ADAR2 in Adar2 KO mice because the same phenomenon was also observed in the cerebral cortex of Adar1 KI mice. Instead, this upregulation was largely observed in sites that were edited by less than 40% in WT mice, regardless of the tissues examined (Fig. 2C–F). These findings suggest that the non-dominant ADAR has little effect on RNA editing for sites that are efficiently edited by the dominant ADAR, whereas non-dominant ADAR seems to negatively affect dominant ADAR-mediated editing for the remaining sites via competition. Although these antagonizing effects have been proposed for inactive ADAR3, and have been reported mainly in REs and miRNAs for ADAR1 and ADAR2 (Chen et al. 2000; Kawahara et al. 2007b; Riedmann et al. 2008; Vesely et al. 2014; Tan et al. 2017), such data indicate that these are not rare events for ADAR1 and ADAR2 even in CDS.

**FIGURE 2.**
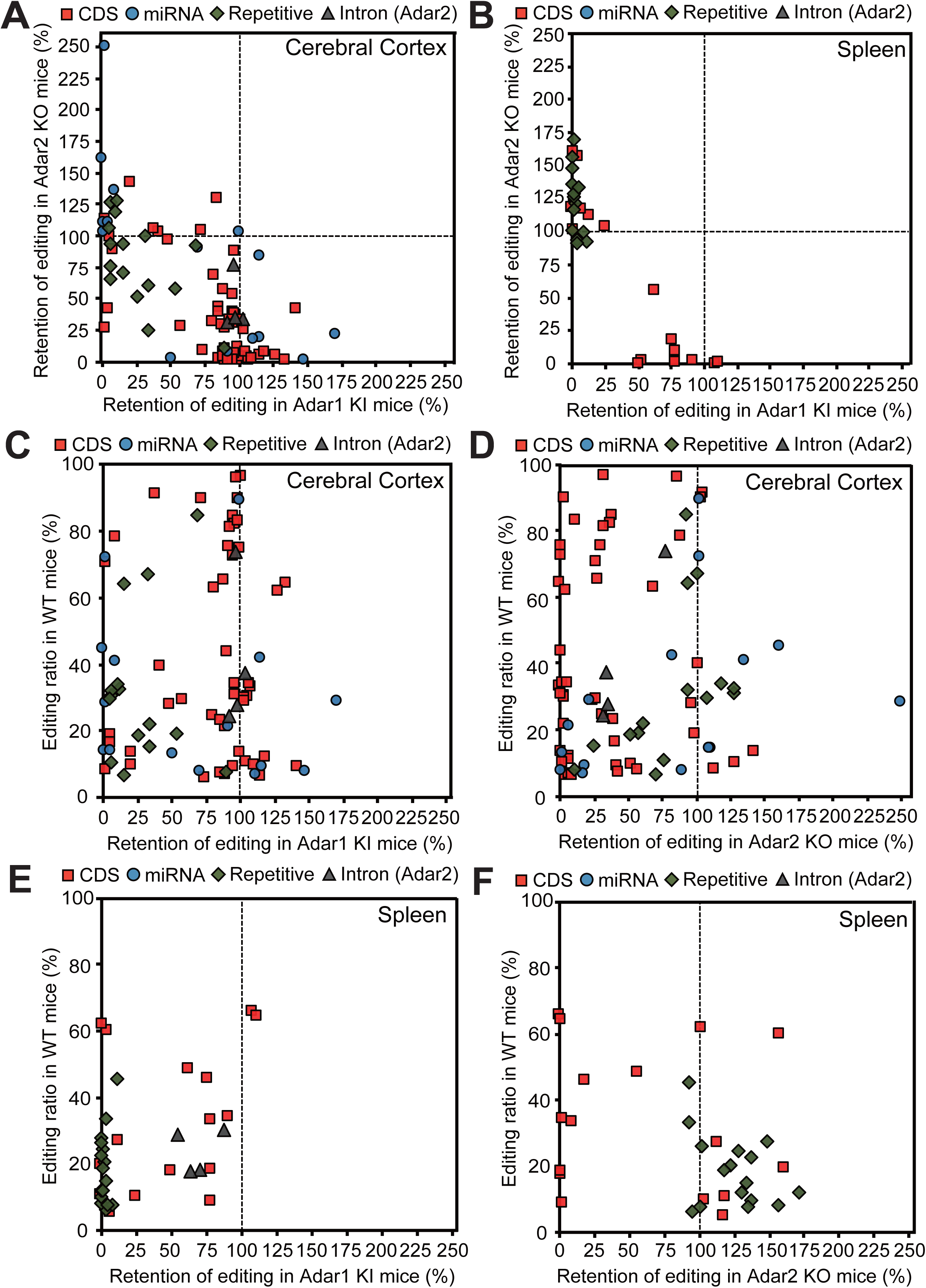
Comparison of editing retention between Adar1 KI and Adar2 KO mice. (*A-B*) The retention of editing in the cerebral cortex (*A*) and spleen (*B*) was compared between *Adar1^E861A/E861A^Ifih^−/−^* mice (Adar1 KI) and *Adar2^−/−^Gria2^R/R^* (Adar2 KO) mice. After calculating the mean editing ratio at each site as the mean of values from three mice for each mutant mouse strain, the retention was calculated by dividing the mean editing ratio of each mutant mouse strain by that of wild-type (WT) mice and shown as a percentage. (*C–F*) Values for the retention of editing in cerebral cortexes isolated from Adar1 KI (*C*) and Adar2 KO (*D*) mice and that in spleens isolated from Adar1 KI (*E*) and Adar2 KO (*F*) mice are displayed. The mean editing ratios of WT mice are displayed on the vertical axis. The red squares, blue circles, green diamonds and grey triangle dots represent editing sites in coding sequences (CDS), microRNAs (miRNAs), repetitive elements and introns, respectively.

### Editing is regulated in a site-specific manner

To know the degree of contribution of each ADAR on each editing site, we calculated the ADAR dominancy, in which 0% dominancy indicated an equal contribution of both ADARs, while 100% dominancy indicated only a single ADAR contribution to the editing of a certain site (Materials and Methods). This analysis demonstrated that the majority of sites in the CDS were dominantly edited by ADAR2, which included Kv1.1 I/V, FLNA Q/R, and CYFIP2 K/E, as well as many sites in glutamate receptor subunits (Fig. 1C, Fig. 3A-B, Supplemental Fig. S2A-B, and Supplemental Table S4), as previously reported (Bhalla et al. 2004; Nishimoto et al. 2008; Riedmann et al. 2008; Stulić and Jantsch 2013). In addition, we showed for the first time that some sites, such as UNC80 S/G, mGluR4 Q/R, NOVA1 S/G, TMEM63B Q/R and SPEG E/G were ADAR2 sites (Supplemental Fig. S3A–D). ADAR2 also contributed dominantly to editing at the SPEG S/G site, especially in the spleen (Supplemental Fig. S3E). In contrast, a certain number of sites in the CDS were ADAR1-dependent, such as three sites in BLCAP and NEIL1 K/R, in addition to AZIN1 E/G and S/G sites (Fig. 1C, Fig. 3A-B, Fig. 4A-B, and Supplemental Table S4), as previously reported (Nishimoto et al. 2008; Riedmann et al. 2008; Yeo et al. 2010; Chen et al. 2013). Furthermore, we identified that UBE2O S/G and DACT3 R/G were edited mainly by ADAR1 (Fig. 4C-D). The CDK13 Q/R site was reported to be edited by ADAR2 (Terajima et al. 2016), whereas the current analysis demonstrated that ADAR1 was a dominant contributor for this site (Fig. 4E). Interestingly, the editing ratios for BLCAP Y/C, DACT3 R/G and CDK13 Q/R sites in the cerebral cortex were higher than those in the spleen where ADAR1 p150 was highly expressed (Fig. 4A, D and E). This suggests that nuclear ADAR1 p110 is the main contributor to this editing, given that their ECS are usually located in the adjacent intron (Levanon et al. 2005). Of note, although the deletion of ADAR2 did not affect editing of these sites in the cerebral cortex, we observed a significant level of RNA editing in Adar1 KI mice, suggesting that ADAR2 edits these sites only in the absence of ADAR1 activity. We further found unique ADAR-dependency in the GABRA3 I/M site. ADAR1 edited the GABRA3 I/M site only in the absence of ADAR2 activity in the cerebral cortex (Fig. 5A). However, both ADAR1 and ADAR2 nearly contributed to this editing in the spleen (Fig. 3B and 5A), which may be due to the difference in dosage of ADAR2 between the two tissues.

**FIGURE 3.**
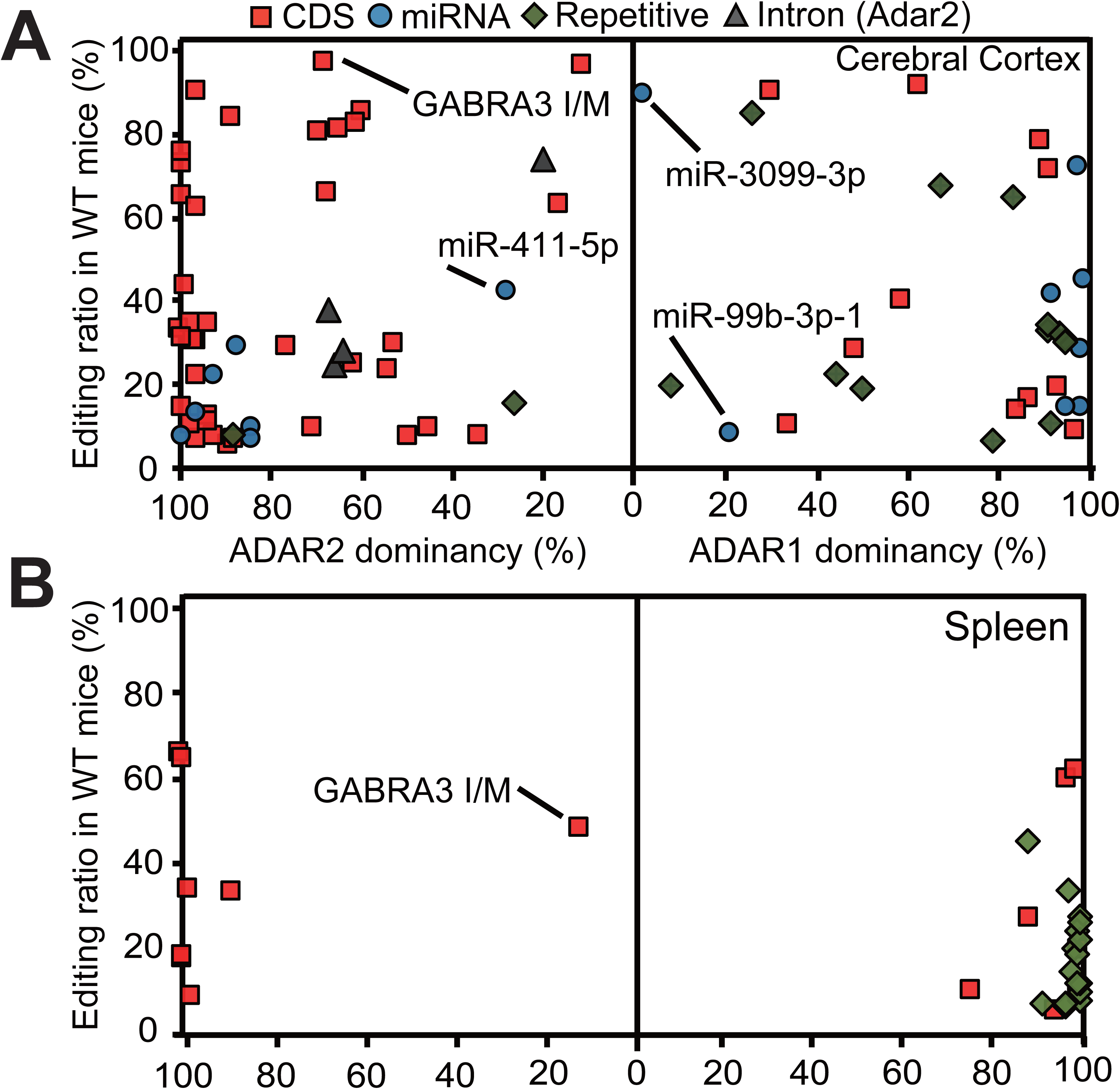
Contribution of each ADAR to each editing site. (*A-B*) ADAR dominancy in the cerebral cortex (*A*) and spleen (*B*). We compared values of editing retention for each site between *Adar1^E861A/E861A^Ifih^−/−^* mice (Adar1 KI) and *Adar2^−/−^Gria2^R/R^* (Adar2) KO mice and defined small values as “A” and large ones as “B”. We calculated ADAR dominancy using the following formula: 100−100×(A/B). Editing ratios in WT mice are shown on the vertical axis. In this figure, 0% and 100% of ADAR dominancy indicates an equal contribution of both ADARs and the sole contribution of a single ADAR to RNA editing at a certain site, respectively. The red squares, blue circles, green diamonds and grey triangle dots represent editing sites in coding sequences (CDS), microRNAs (miRNAs), repetitive elements and introns, respectively.

**FIGURE 4.**
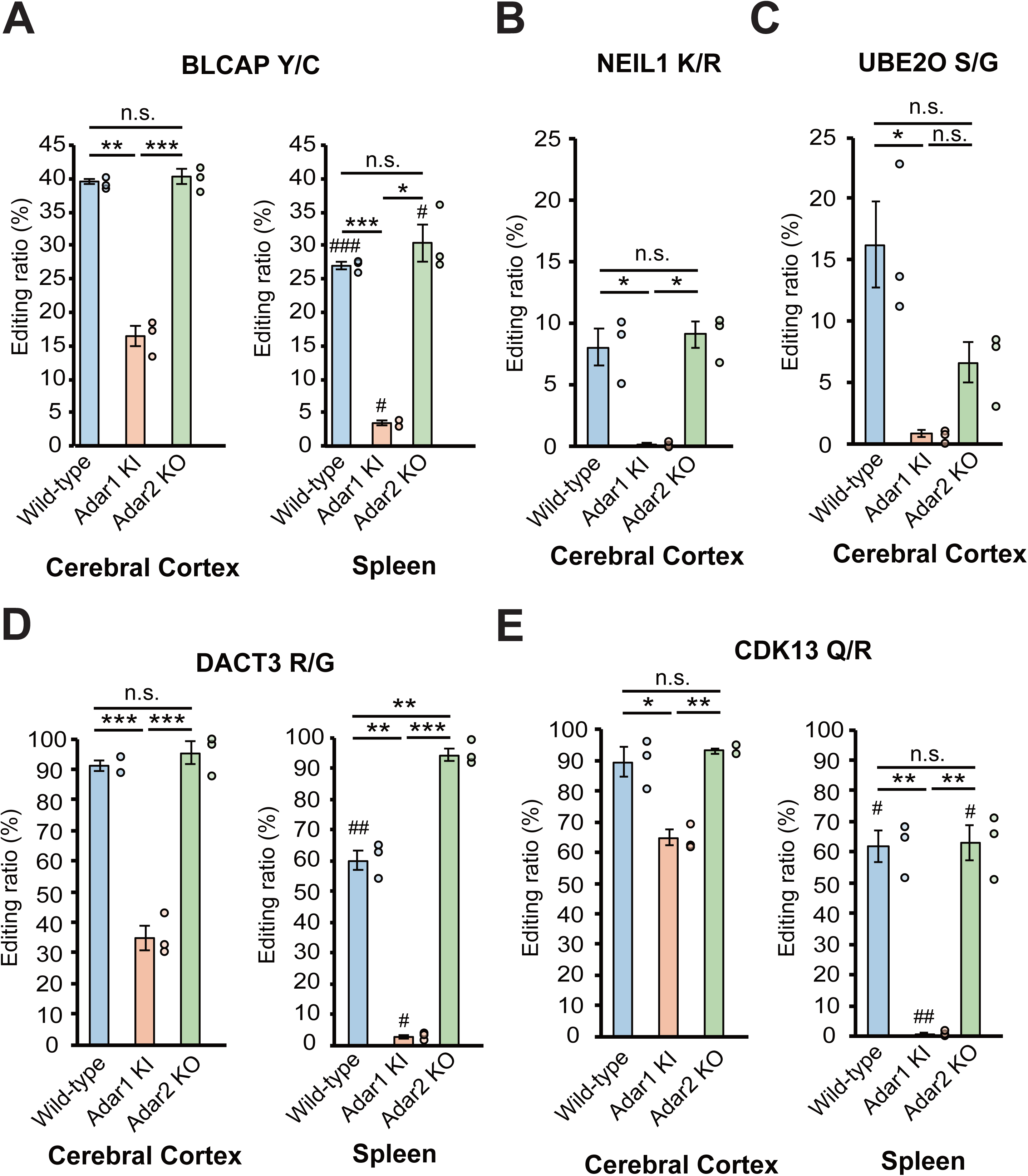
Retention of RNA editing at known and novel ADAR1 sites in Adar1 KI and Adar2 KO mice. (*A–E*) Editing ratios for the BLCAP tyrosine/cysteine (Y/C) site (*A*), NEIL1 lysine/arginine (K/R) site (*B*), UBE2O serine/glycine (S/G) site (*C*), DACT3 arginine/glycine (R/G) site (*D*) and CDK13 glutamine/arginine (Q/R) site (*E*) are shown. Editing ratios are displayed as the mean ± SEM (n=3 mice for each group; Student’s *t*-test, \**p* < 0.05, \*\**p* < 0.01, \*\*\**p* < 0.001, n.s., not significant). The editing ratio of each mouse is also displayed as a circle on the right side of each column. Significant differences in editing ratios between the cerebral cortex and spleen in the same mutant mice are indicated by hashes (^#^*p* < 0.05, ^##^*p* < 0.01, ^###^*p* < 0.001). Adar1 KI, *Adar1^E861A/E861A^Ifih^−/−^* mice; Adar2, *Adar2^−/−^Gria2^R/R^*.

**FIGURE 5.**
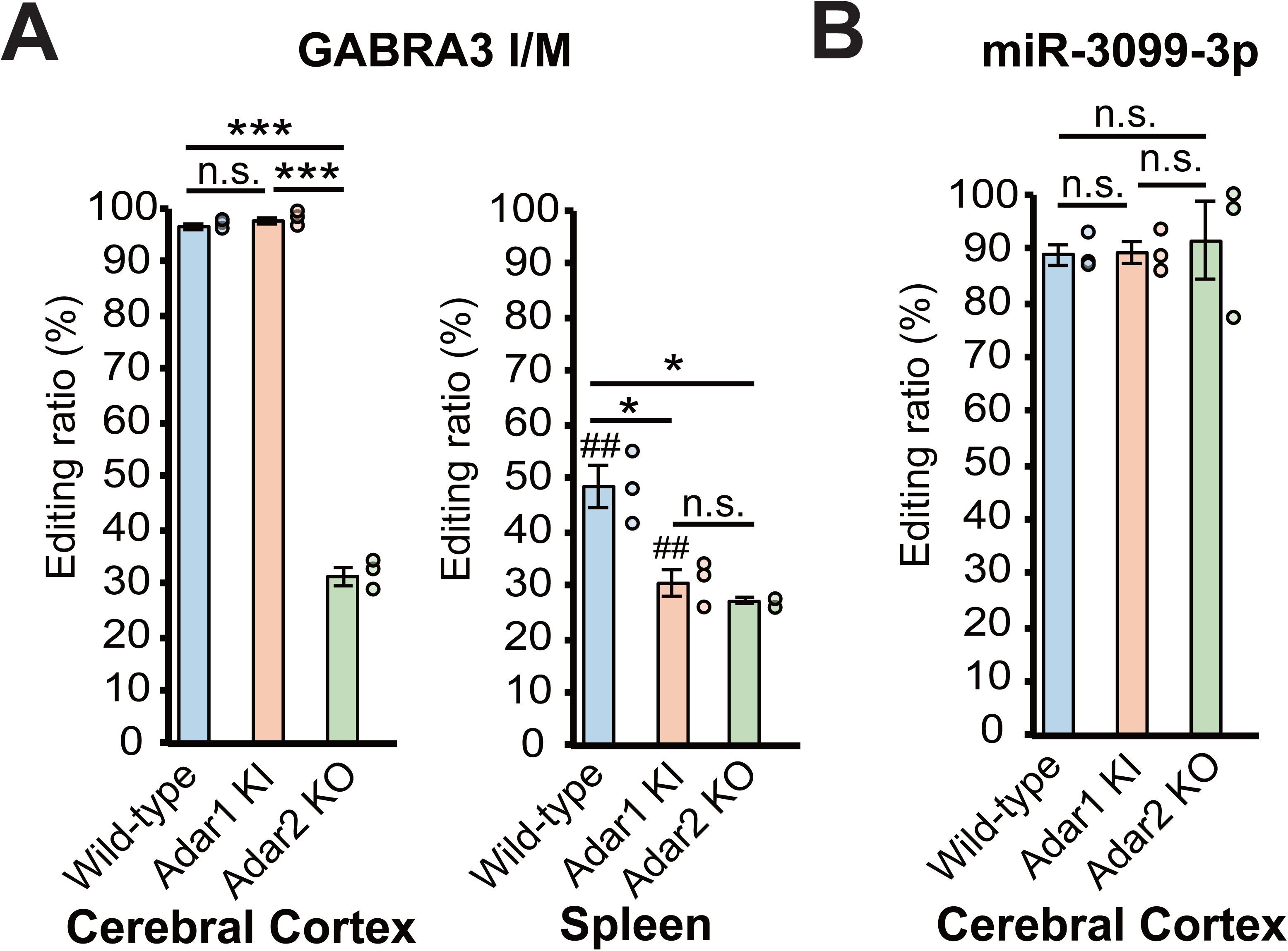
Site-specific regulation of RNA editing. (*A-B*) Editing ratios for the GABRA3 isoleucine/methionine (I/M) site (*A*) and the site in miR-3099-3p (*B*) in indicated tissues isolated from wild-type (WT), *Adar1^E861A/E861A^Ifih^−/−^* mice (Adar1 KI) and *Adar2^−/−^Gria2^R/R^* (Adar2 KO) mice are shown. Editing ratios are displayed as the mean ± SEM (n=3 mice for each group; Student’s *t*-test, \**p* < 0.05, \*\*\**p* < 0.001, n.s., not significant). The editing ratio of each mouse is also displayed as a circle on the right side of each column. Significant differences in editing ratios between the cerebral cortex and spleen in the same mutant mice are indicated by hashes (^##^*p* < 0.01).

Regarding miRNA editing, we reconfirmed that sites in miR-423-5p, miR-376b-3p, miR-376c-3p, miR-151-3p and miR-99b-3p were dominantly edited by ADAR1, whereas ADAR2 was the dominant enzyme for sites in miR-27a-5p, miR-376a-2-5p, miR-379-5p, miR-381-3p and miR-99a-5p (Fig. 3A-B, and Supplemental Table S4) (Kawahara et al. 2007a; Kawahara et al. 2007b; Kawahara et al. 2008b; Vesely et al. 2014; Nishikura 2016). In addition, for the first time, this analysis identified ADAR2 as the dominant enzyme in the editing of miR-27b-5p. It is noteworthy that most sites in the 3p-strand were edited by ADAR1, whereas ADAR2 was responsible for editing most of the sites in the 5p-strand, suggesting differences in accessibility between the two enzymes. Interestingly, miR-3099-3p was ∼90% edited in the cerebral cortex, with an editing ratio preserved in both ADAR1 KI and ADAR2 KO mice (Fig. 5B). Although this site was previously reported as a possible ADAR1 site (Vesely et al. 2014), this result suggests that this is the first case in which a highly edited site shows no preference for a specific ADAR (Fig. 3A). A similar phenomenon was also observed for miR-411-5p and miR-99b-3p-1 (Fig. 3A, Supplemental Fig. S4A-B and Supplemental Table S4), which may be the reason why the ADAR responsible for editing miR-411-5p was found not to be identical among past studies (Kawahara et al. 2008b; Vesely et al. 2014; Nishikura 2016). Taken together, the editing of miR-3099-3p and miR-411-5p may be used as control substrates to analyze the difference in properties between ADAR1 and ADAR2 *in vivo* and *in vitro*.

Next, we investigated the similarity of ADAR dominancy between the cerebral cortex and spleen. This analysis demonstrated a high correlation for most sites in CDS and miRNAs (Fig. 6A–C). For instance, we observed high ADAR1 dependency for three sites, BLCAP, CKD13 Q/R and DACT3 R/G, in both the cerebral cortex and spleen (Fig. 6A-B). In contrast, although both ADAR1 and ADAR2 participated in the editing of REs in the cerebral cortex, ADAR1 was the dominant enzyme in the spleen where the presence of ADAR2 had a negative effect (Fig. 3A-B and Fig. 6A-B). As a result, we observed differences between the cerebral cortex and spleen with regard to the ADAR responsible for several editing sites in REs (Fig. 6C).

**FIGURE 6.**
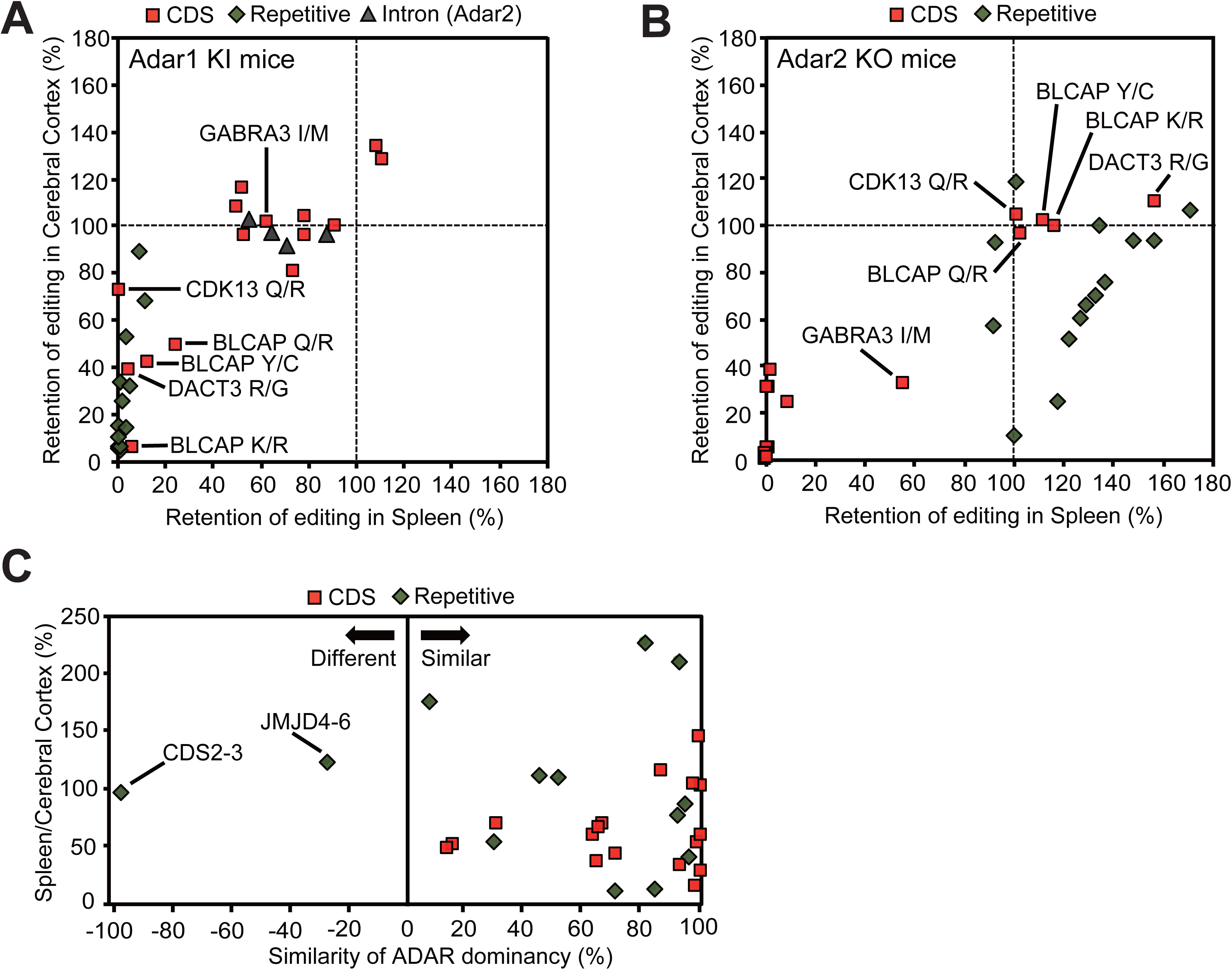
Comparison of editing retention between the cerebral cortex and spleen. (*A-B*) The retention of editing in *Adar1^E861A/E861A^Ifih^−/−^* mice (Adar1 KI) mice (*A*) and *Adar2^−/−^Gria2^R/R^*(Adar2 KO) mice (*B*) was compared between the cerebral cortex and spleen. (*C*) Degree of similarity of ADAR dominancy between the cerebral cortex and spleen. The value of ADAR dominancy was compared between the cerebral cortex and spleen; small values were defined as “C” and large ones as “D”; the similarity of ADAR dominancy was calculated using the following formula: 100×(C/D). In cases where ADAR1 is dominant in a certain tissue and ADAR2 is dominant in another tissue, this is shown as a negative value to express the difference in ADAR dominancy. In this figure, 100% and −100% on the horizontal axis indicates the contribution of the same and different ADARs, respectively, to a certain editing site in the cerebral cortex and spleen. The relative value calculated by dividing the editing ratio in the spleen by that in the cerebral cortex isolated from wild-type (WT) mice is shown on the vertical axis as a percentage. The red squares, green diamonds and grey triangle dots represent editing sites in coding sequences (CDS), repetitive elements and introns, respectively.

### Some sites require interplay between ADAR1 and ADAR2 for efficient RNA editing

It has been reported that among the five editing sites of brain-specific 5-HT_2C_R encoded by the *Htr2c* gene, the A and B sites are dominantly edited by ADAR1, while the E, C and D sites are preferentially edited by ADAR2 (Burns et al. 1997; Liu et al. 1999; Nishikura 2016), as confirmed by this study (Fig. 7A). However, although editing at the B site almost disappeared in Adar1 KI mice, the deletion of ADAR2 also reduced the editing ratio significantly from 71% to 19% (Fig. 7A). This result suggests that coordinated interplay between ADAR1 and ADAR2, in which preceding editing by ADAR2 might alter the secondary structure, was required for efficient editing at this site. Indeed, the editing ratio of the B site in the rat brain was higher than that obtained by *in vitro* RNA editing assay with recombinant (r)ADAR1 (Liu et al. 1999). However, the B site editing was not increased by a simple combination of rADAR1 and rADAR2 (Chen et al. 2000). In addition, although subtle or non-significant, similar phenomena were also observed at the UBE2O S/G site in the cerebral cortex (ADAR1 site), the CACNA1D I/M site in the cerebral cortex and the FLNB Q/R site in the spleen (ADAR2 sites), all of which do not have additional editing sites within the same dsRNA structure targeted by a different ADAR (Fig. 4C and Fig. 7B-C). Nevertheless, given that inactive ADAR1 is expressed in ADAR1 KI mice, these results suggest that the editing activity, but not the binding capacity, of non-dominant ADAR is critical for promoting RNA editing of these sites. Intriguingly, preceding editing of the intronic F site of 5-HT_2C_R altered the editing pattern of some exonic sites (Flomen et al. 2004). Therefore, although the mechanism underlying coordinated interplay between ADAR1 and ADAR2 remains currently unknown, one possibility is that the non-dominant ADAR has an additional editing site in the intronic ECS and that preceding editing of this site may alter the secondary structure, leading to efficient editing by the dominant ADAR.

**FIGURE 7.**
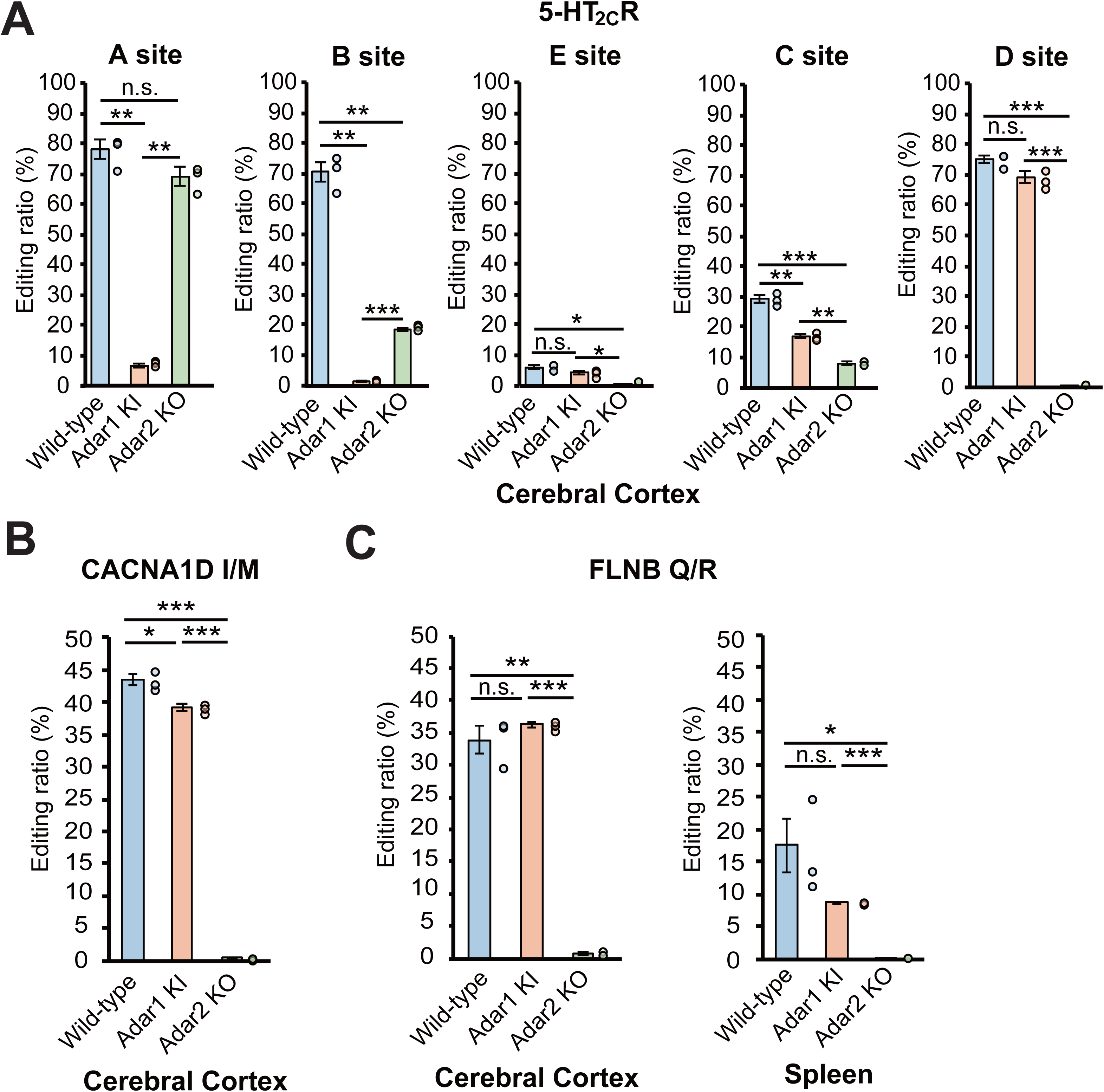
Cases that require non-dominant ADAR for efficient RNA editing. (*A–C*) Editing ratios of five sites in the serotonin 5-HT_2C_R (*A*), CACNA1D isoleucine/methionine (I/M) site (*B*) and FLNB glutamine/arginine (Q/R) site (C) in the indicated tissues isolated from wild-type (WT), *Adar1^E861A/E861A^Ifih^−/−^* mice (Adar1 KI) and *Adar2^−/−^Gria2^R/R^* (Adar2 KO) mice are shown. Editing ratios are displayed as the mean ± SEM (n=3 mice for each group; Student’s *t*-test, \**p* < 0.05, \*\**p* < 0.01, \*\*\**p* < 0.001, n.s., not significant). The editing ratio of each mouse is also displayed as a circle on the right side of each column.

We further analyzed the combination of editing within the same transcripts. As expected, the majority of *Htr2c* transcripts were edited only at the D site, with or without the C site, in Adar1 KI mice (Fig. 8A). In contrast, transcripts edited at the A site, with or without the B site, occupied more than 60% in Adar2 KO mice (Fig. 8A). Given that the two sites in CACNA1D are dominantly edited by ADAR2, no substantial alteration of the editing combination at the two sites was observed in Adar1 KI mice, whereas only unedited transcripts were observed in Adar2 KO mice (Fig. 8B). Conversely, no substantial alteration of the editing combination at the three sites in BLCAP was observed in the spleen of Adar2 KO mice, whereas more than 90% of transcripts were unedited in Adar1 KI mice (Fig. 8C). Intriguingly, ADAR2 edited either Y/C or Q/R sites, or both to some extent but not the K/R site, which was the lowest edited site of BLCAP in the cerebral cortex of Adar1 KI mice, and therefore the combination pattern was different from that in the spleen. However, except for *Htr2c* transcripts, the multiple editing sites within the same dsRNA structure are usually targeted by one dominant ADAR. This suggests that the preferred dsRNA structure may differ for ADAR1 and ADAR2, which requires further investigation.

**FIGURE 8.**
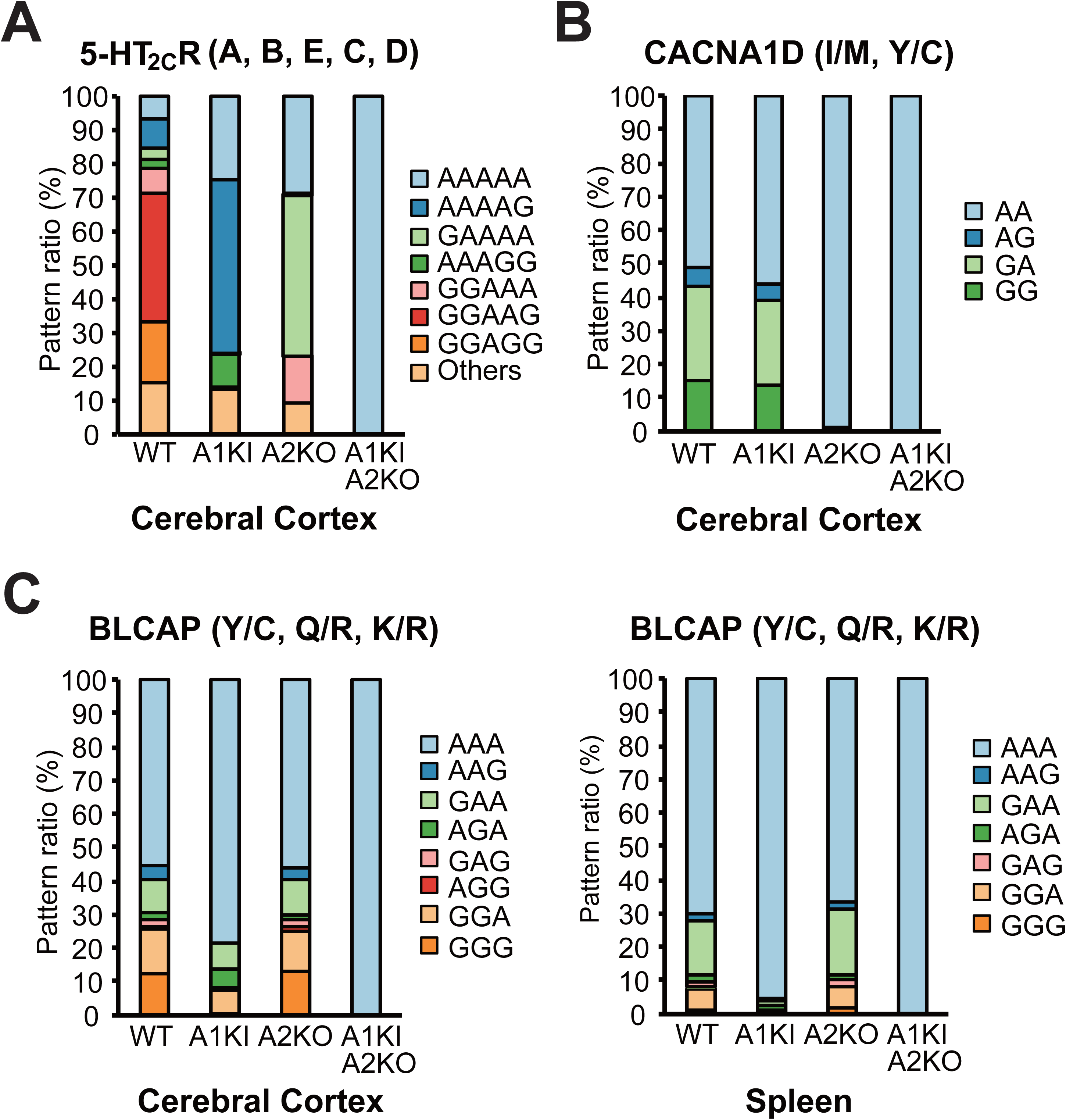
Comparison of the combined editing pattern in the same transcripts among Adar mutant mice. (*A–C*) The frequency of each combined editing pattern of the five sites (A, B, E, C and D in order) in the serotonin 5-HT_2C_R (*A*), the two sites (isoleucine/methionine [I/M] and tyrosine/cysteine [Y/C] in order) in CACNA1D (*B*) and the three sites (Y/C, glutamine/arginine [Q/R] and lysine/arginine [K/R] in order) in BLCAP (*C*) in the indicated tissues isolated from wild-type (WT; n = 3 mice), *Adar1^E861A/E861A^Ifih^−/−^* mice (Adar1 KI; n = 3 mice), *Adar2^−/−^Gria2^R/R^* (Adar2 KO; n = 3 mice) and Adar1 KI Adar2 KO (n = 2 mice) mice is displayed as a percentage.

### Adar1 KI Adar2 KO mice are viable with a complete absence of editing

Adar2 KO (*Adar2^−/−^Gria2^R/R^*) mice survive until adulthood (Higuchi et al. 1993). In addition, although *Adar1^E861A/E861A^* mice show embryonic lethality, Adar1 (*Adar1^E861A/E861A^Ifih^−/−^*) KI mice can survive with a normal life-span (Liddicoat et al. 2015; Heraud-Farlow et al. 2017). However, given that ADAR1-mediated RNA editing in CDS is independent of MDA5 activation and that the non-dominant ADAR can edit many sites in CDS and miRNAs in the absence of dominant ADAR1 activity in compensation, we generated Adar1 KI Adar2 KO mice to examine whether RNA editing completely disappeared and to observe its phenotypic consequences. We found that Adar1 KI Adar2 KO mice were viable until adulthood and showed a significantly smaller body size compared to WT and Adar2 KO mice (Fig. 9A). However, this small body size was comparable to that of Adar1 KI mice, which is known to be small (Liddicoat et al. 2015; Heraud-Farlow et al. 2017). Mating of Adar1 KI Adar2 KO mice was difficult because of their small sizes; therefore, we successfully obtained pups from both male and female Adar1 KI Adar2 KO mice by *in vitro* fertilization, which indicated that Adar1 KI Adar2 KO mice were fertile. Therefore, although we could not exclude the possibility that additional abnormalities may have been detected by undertaking more detailed examinations, as reported in Adar2 KO mice that showed a myriad of subtle phenotypes (Horsch et al. 2011), a cumulative effect on critical phenotypes due to the inactivation of ADAR1 and the deletion of ADAR2 in mice was not apparent.

**FIGURE 9.**
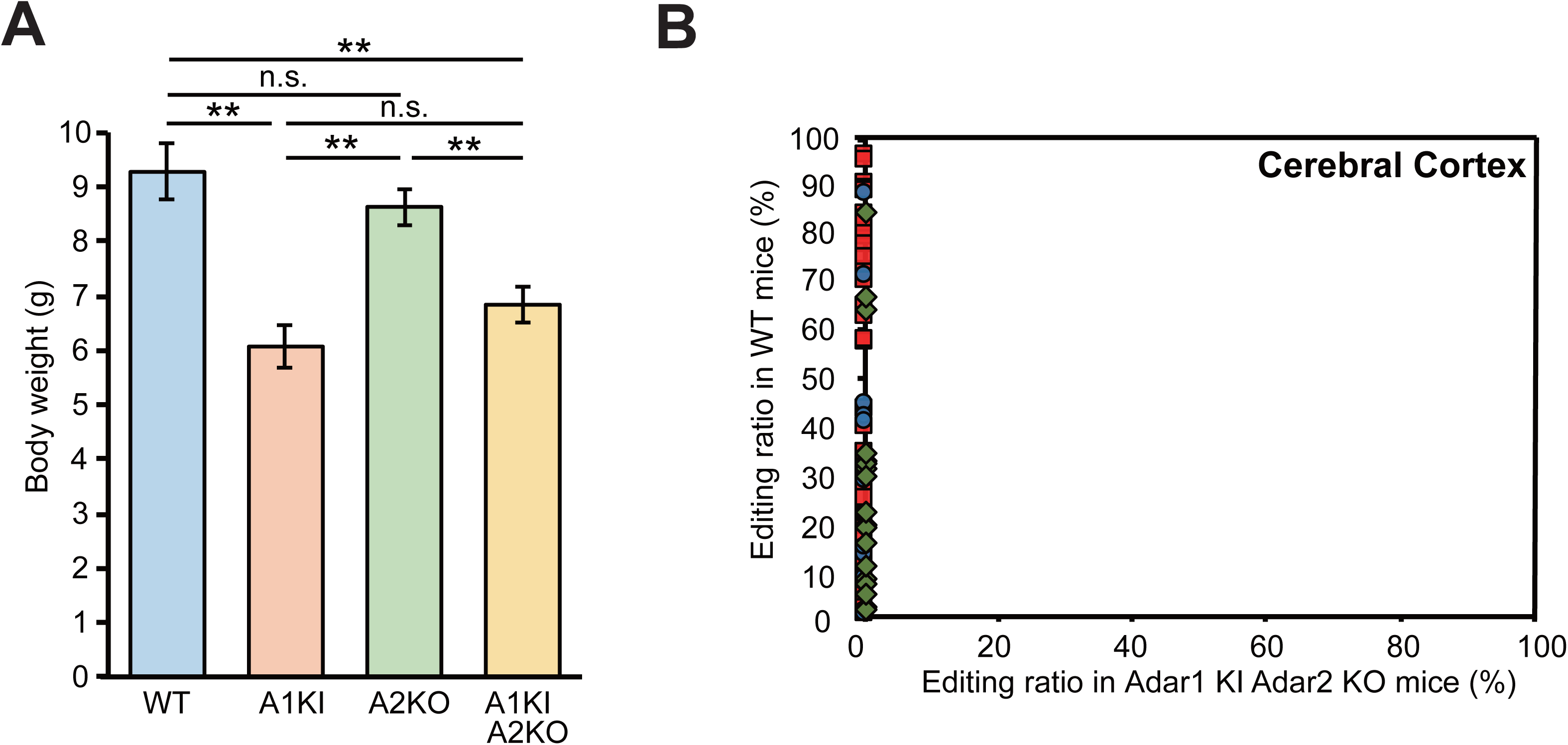
Complete absence of RNA editing in Adar1 KI Adar2 KO mice. (*A*) The body weights of wild-type (WT; n = 4 mice), *Adar1^E861A/E861A^Ifih^−/−^* mice (Adar1 KI; n = 6 mice), *Adar2^−/−^Gria2^R/R^* (Adar2 KO; n = 9 mice) and Adar1 KI Adar2 KO (n = 11 mice) mice at 10 days of age are displayed as the mean ± SEM (Student’s *t*-test, \*\**p* < 0.01, n.s., not significant). (*B*) Editing ratios of all sites examined in the cerebral cortex were compared between WT and Adar1 KI Adar2 KO mice. Values are displayed as the mean of the values of three WT and two Adar1 KI Adar2 KO mice. The red squares, blue circles, green diamonds and grey triangle dots represent editing sites in coding sequences (CDS), microRNAs (miRNAs), repetitive elements and introns, respectively.

Finally, using Adar1 KI Adar2 KO mice, we analyzed the editing ratios of all sites examined in Adar1 KI and Adar2 KO mice. This analysis demonstrated a complete loss of RNA editing in both the cerebral cortex and spleen of Adar1 KI Adar2 KO mice (Fig. 9B). Using total RNA-sequencing (RNA-seq) analysis, we further examined whether editing at certain sites was maintained in these mutant mice. However, although more than 3,000 known editing sites were detected in WT mice, editing sites were not detected in Adar1 KI Adar2 KO mice except for the GluR2 Q/R site, which was knocked-in at the genomic DNA level (Supplemental Table S5). These results suggest that ADAR1 and ADAR2 are the enzymes responsible for all editing sites *in vivo* and that critical editing sites differ between ADAR1 and ADAR2.

In summary, we have provided comprehensive and quantitative data for RNA editing in all sites that are known to be conserved or may be conserved in the CDS and miRNAs of WT, Adar1 KI, Adar2 KO, and Adar1 KI Adar2 KO mice. This will be an invaluable resource for furthering our understanding of which Adar dominantly contributes to the editing of each site. In addition, total RNA-seq data obtained from Adar1 KI Adar2 KO mice can contribute to excluding false-positive sites. We have demonstrated that editing is regulated by a site-specific mechanism related to the interplay between ADAR1 and ADAR2, which sometimes cooperatively edited certain sites; however, the non-dominant ADAR frequently had an inhibitory role on editing by the dominant ADAR. These findings were obtained by comparing the editing ratios of each specific site between WT, Adar1 KI and Adar2 KO mice. Although RNA editing at each site is affected by many factors, including secondary structure and the presence of neighboring and opposing nucleotides and regulators such as aminoacyl tRNA synthetase complex-interacting multifunctional protein 2 (AIMP2) (Riedmann et al. 2008; Kuttan and Bass 2012; Tan et al. 2017; Wang et al. 2018), our study showed that the tissue-specific relative dosage of each ADAR is a factor underlying the tight regulation of site-specific RNA editing.

## MATERIAL AND METHODS

### Mouse administration

Mice were maintained on a 12-h/12-h light/dark cycle at a temperature of 23 ± 1.5°C with a humidity of 45 ± 15% as previously described (Nakahama et al. 2018). All experimental procedures that involved mice were performed in accordance with protocols approved by the Institutional Animal Care and Use committee of Osaka University.

### Mutant mice

Cryopreserved spermatozoa of Adar2 KO mice were obtained from the Mutant Mouse Resource and Research Center (RRID: MMRRC_034679-UNC) and *in vitro* fertilization was performed at the Institute of Experimental Animal Sciences Faculty of Medicine, Osaka University. *Adar1^E861A/+^* mice that harbor a heterozygous editing-inactive E861A point mutation were generated by genome editing using the CRISPR/Cas9 system at The Genome Editing Research and Development Center, Graduate School of Medicine, Osaka University. Briefly, pronuclear-stage mouse embryos (CLEA Japan Inc., Tokyo, Japan) were electroporated to introduce *Cas9* mRNA, single guide RNA (sgRNA: TCCGGGAGATGATTTCGGCA) and single-stranded donor oligonucleotides (ssODN: ATGGCTGTGCCCATCTTGCTTACCTGATGAAGCCCCTCCGGGAGATGATT<u>G</u>CGGCATG GCAGTCATTGACCGTCTCTCCCTTCAGGCTGAGAGAGTCCCCTTT); these introduced a point mutation at the corresponding codon (underlined G in the target nucleotide). Mouse embryos that developed to the two-cell stage were transferred into the oviducts of female surrogates. *Adar1^E861A/E861A^* mice were then established by crossing with *Adar1^E861A/+^* mice. We did not find a difference in phenotypes between the obtained *Adar1^E861A/E861A^* mice and *Adar1^E861A/E861A^* mice that were previously established by a conventional method (Liddicoat et al. 2015; Nakahama et al. 2018). Furthermore, Adar1 KI mice were obtained by crossing *Adar1^E861A/+^* mice with *Ifih^−/−^* mice as reported previously (Nakahama et al. 2018). To establish Adar1 KI Adar2 KO mice, we generated *Adar1^E861A/+^Ifih^−/−^Adar2^−/−^Gria2^R/R^* mice. During this procedure, repeated backcrossing was required to induce homologous recombination, given that the *Adar1* and *Gria2* genes localize to the same chromosome. Finally, we performed *in vitro* fertilization using sperm and ova collected from *Adar1^E861A/+^Ifih^−/−^Adar2^−/−^Gria2^R/R^* mice. Genotyping of the *Adar1* gene was performed by direct Sanger sequencing of the PCR products amplified from the region, including the point mutation. All mice used in experiments were in a C57BL/6J background.

### Western blot analysis

Tissue lysates from mouse cerebral cortex and spleen were prepared and stored at −80°C until use as described previously (Miyake et al. 2016). Lysates were then separated using sodium dodecyl sulfate (SDS)-polyacrylamide gel electrophoresis (PAGE), transferred to a polyvinylidene difluoride (PVDF) membrane (Bio-Rad, Hercules, CA, USA) and immunoblotted with primary antibodies using a SNAP i.d^®^ 2.0 Protein Detection System (Merck Millipore, Burlington, MA, USA) as previously described (Nakahama et al. 2018). The primary antibodies used were as follows: mouse monoclonal anti-ADAR1 antibody (15.8.6; Santa Cruz Biotechnology, Dallas, TX, USA), mouse monoclonal anti-ADAR2 antibody (1.3.1; Santa Cruz Biotechnology) and mouse monoclonal anti-GAPDH (M171-3; MBL).

### Total RNA preparation

Total RNA was extracted from the cerebral cortex and spleen collected from eight-week-old male mice using TRIzol reagent (Thermo Fisher Scientific, Waltham, MA, USA) in accordance with the manufacturer’s protocol. The RNA concentration was measured using a NanoDrop One (Thermo Fisher Scientific) and stored at −80°C until use after adjustment to 1 µg/µL.

### Preparation of Ion amplicon libraries for RNA editing sites

After 1 µg of total RNA from each tissue was treated with DNase I (Thermo Fisher Scientific) at 37°C for 20 min, cDNA was synthesized by reverse-transcription (RT) using a SuperScript™ III First-Strand Synthesis System (Thermo Fisher Scientific) according to the manufacturer’s instructions (Fig. 1A). Random hexamers were used as RT primers for editing sites in introns and miRNAs, while oligo(dT) primers were used for sites in mRNAs to avoid possible contamination of pre-mRNA fragments. A first round of PCR was performed using cDNA, Phusion Hot Start High-Fidelity DNA Polymerase (Thermo Fisher Scientific) and first primers that were editing-site specific (Fig. 1A and Supplemental Table S6). A second round of PCR was then performed using an aliquot of the first PCR product as a template and second primers that were editing-site specific; an A adaptor (5’-CCATCTCATCCCTGCGTGTCTCCGACTCAG-3’), an Ion Xpress Barcode™ and a trP1 adaptor (5’-CCTCTCTATGGGCAGTCGGTGAT-3’) were in forward and reverse primers, respectively (Fig. 1A and Supplemental Table S6). All second PCR products were designed to be 190 to 200 bp in length. After gel purification, the concentration of each PCR product was measured using a NanoDrop One and then equal amounts of 50–300 PCR products were combined. After a quality check using a 2100 Bioanalyzer (Agilent Technologies, Santa Clara, CA, USA) with a High Sensitivity DNA kit, the resultant amplicon library samples were subjected to deep sequencing using an Ion Torrent™ Personal Genome Machine™ (Ion PGM) system (Thermo Fisher Scientific) at the CoMIT Omics Center, Graduate School of Medicine, Osaka University.

### Quantification of the RNA editing ratio with Ion amplicon sequencing reads

An RNA editing ratio for each site was calculated with its read data generated by an Ion PGM. For each amplicon sequence, the prefix of length k, which we termed the k-prefix, was taken and compared with the k-prefix of a read generated by the sequencer. Note that the k-prefix of an amplicon sequence was derived from a specific primer sequence within the amplicon, meaning that the k-prefixes derived from all amplicon sequences were unique. Also, k was set to six in this study. If the k-prefix of the read was identical to that of the amplicon, it was investigated to determine whether a known editing site in the amplicon was edited. This was repeated until all sequence reads were scanned. Using this simple process, implemented with an in-house script, we calculated the editing ratio by dividing the number of edited reads by that of the total reads for each site. We set the minimum threshold for the number of total reads to 1,000. Then, the mean editing ratio at each site was calculated using the editing ratios obtained from three WT, three Adar1 KI, three Adar2 KO and two Adar1 KI Adar2 KO mice.

### Calculation of editing retention

To determine how much editing was retained in tissues from Adar1 KI and Adar2 KO mice, the mean editing ratio of each mutant mouse was divided by that of WT mice to calculate the value for the retention of editing at each site. We only considered sites with more than a 5% editing ratio in WT mice for this analysis.

### Calculation of ADAR dominancy

To quantify to what extent each ADAR is responsible for the editing of each site, we compared the value of the editing retention for each site between Adar1 KI and Adar2 KO mice, and defined the small value as “A” and the large one as “B”. We calculated ADAR dominancy using the following formula: 100−100×(A/B). In this calculation, 0% and 100% indicate an equal contribution of both ADARs and a dominant contribution of a single ADAR to the RNA editing of a certain site, respectively. We only considered sites with more than a 5% editing ratio in WT mice for this analysis.

### Similarity of ADAR dominancy between tissues

To express a similarity or difference of ADAR dominancy between the cerebral cortex and spleen, we compared the value of ADAR dominancy between the cerebral cortex and spleen, and defined the small value as “C” and the large one as “D”. We calculated the similarity of ADAR dominancy using the following formula: 100×(C/D). In cases where ADAR1 is dominant in a certain tissue and ADAR2 is dominant in another tissue, this was shown as a negative value. Therefore, 100% indicates the contribution of the same ADAR between the cerebral cortex and spleen to a certain editing site, while −100% indicates the contribution of different ADARs between the cerebral cortex and spleen. We only considered sites with more than a 5% editing ratio in WT mice for this analysis.

### Quantification of the RNA editing ratio by Sanger sequencing

The editing ratio was analysed by Sanger sequencing as described previously with minor modifications (Miyake et al. 2016). In brief, 100 ng of each total RNA was incubated with 0.1 U/µl DNase I (Thermo Fisher Scientific) at 37°C for 20 min and then the denatured RNAs were reverse transcribed into cDNAs using the SuperScript™ III First-Strand Synthesis System (Thermo Fisher Scientific) with random hexamers. PCR was performed with Phusion Hot Start High-Fidelity DNA Polymerase (Thermo Fisher Scientific) and the following primers: Azin1-Fw1 (5’-GATGAGCCAGCCTTCGTGT-3’) and Azin1-Rv1 (5’-TGGTTCGTGGAAAGAATCTGC-3’) for mature mRNA, and Azin1-Fw2 (5’-TGAGACTTATGCCTGATCGTTG-3’) and Azin1-Rv2 (5’-CCAGCAAATCTAAACTGTCACTCA-3’) for pre-mRNA. After gel purification, each RT-PCR product was directly sequenced using the following primer: 5’-CAAGGAAGATGAGCCTCTGTTT-3’. The editing ratio was determined as the % ratio of the “G” peak over the sum of the “G” and “A” peaks of the sequencing chromatogram.

### Total RNA-sequencing analysis

After ribosomal RNAs were removed from total RNA using a Ribo-Zero rRNA Removal Kit (Illumina, San Diego, CA, USA), a strand-specific RNA library was prepared using SureSelect Strand Specific RNA (Agilent) in accordance with the manufacturer’s instructions as previously described (Nakahama et al. 2018). The library samples were then subjected to deep sequencing using an Illumina NovaSeq 6000 with 100-bp paired-end reads at Macrogen (Kyoto, Japan).

### Genome-wide identification of editing sites

We adopted a genome-wide approach to identify editing sites with total RNA-seq reads as previously described (Nakahama et al. 2018) but with modifications. In brief, sequence reads were mapped onto a reference mouse genome (NCBIM37/mm9) with a spliced aligner HISAT2 (Kim et al. 2015). The mapped reads were then processed by adding read groups, and sorting and marking duplicates with the tools AddOrReplaceReadGroups and MarkDuplicates compiled in GATK4 (McKenna et al. 2010). GATK SplitNCigarReads, BaseRecalibrator and ApplyBQSR were used to split ‘n’ trim and reassign mapping qualities, which output analysis-ready reads for the subsequent variant calling. The GATK HaplotypeCaller was run for variant detection, in which the stand-call-conf option was set to 20.0 and the dont-use-soft-clipped-bases option was used. The results of variant calling were further filtered by GATK VariantFiltration using Fisher strand values (FS) > 30.0 and quality by depth values (QD) < 2.0 as recommended by the GATK developer for RNA-seq analysis. The remaining variants that were expected to be of high quality were annotated with ANNOVAR (Wang et al. 2010). Among these variants, we picked up known editing sites registered in RADAR (Ramaswami and Li 2014). Finally, A-to-I editing ratios in each sample were calculated by dividing the allelic depth by the read depth for the editing sites shown in the annotated results.

### Analysis of dsRNA structure

Potential secondary dsRNA structure was calculated using Mfold (Zuker 2003).

### Statistical analyses

A two-tailed Student’s *t*-test was used as indicated in each figure legend. All values are displayed as the mean ± standard error of the mean (SEM). Non-significance is displayed as n.s., while statistical significance is displayed as *p* < 0.05 (* or ^#^), *p* < 0.01 (** or ^##^) or *p* < 0.001 (*** or ^###^).

## DATA DEPOSITION

The RNA-seq data used in this study are available through the DNA Data Bank of Japan (DDBJ) under accession number DRA007927. The mfold web server is open source software for the prediction of nucleic acid folding and hybridization (https://unafold.rna.albany.edu/?q=mfold/RNA-Folding-Form).

## ACKNOWLEDGEMENT

We thank all the staff in the Genome Editing Research and Development Center, the Center for Medical Research and Education and the CoMIT Omics Center, Graduate School of Medicine, Osaka University, for technical support. Computations were partially performed on the NIG supercomputer at ROIS National Institute of Genetics, Japan.

## FUNDING

This work was supported by Grants-in-Aid KAKENHI [17K19352 to Y. Kawahara, 18K15186 and 15K19126 to T.N., 18K11526 and 15K00401 to Y. Kato] from the Ministry of Education, Culture, Sports, Science and Technology (MEXT) of Japan and by grants from SENSHIN Medical Research Foundation, The Mochida Memorial Foundation for Medical and Pharmaceutical Research (to Y. Kawahara), Nagao Memorial Fund (to T.N.), and the Takeda Science Foundation (to Y. Kawahara and T.N.). P.H.C.C. was supported by a MEXT scholarship.

## CONFLIT OF INTEREST

None declared

