## Supplemental Figures for "A comparative analysis among ADAR mutant mice reveals site-specific regulation of RNA editing"

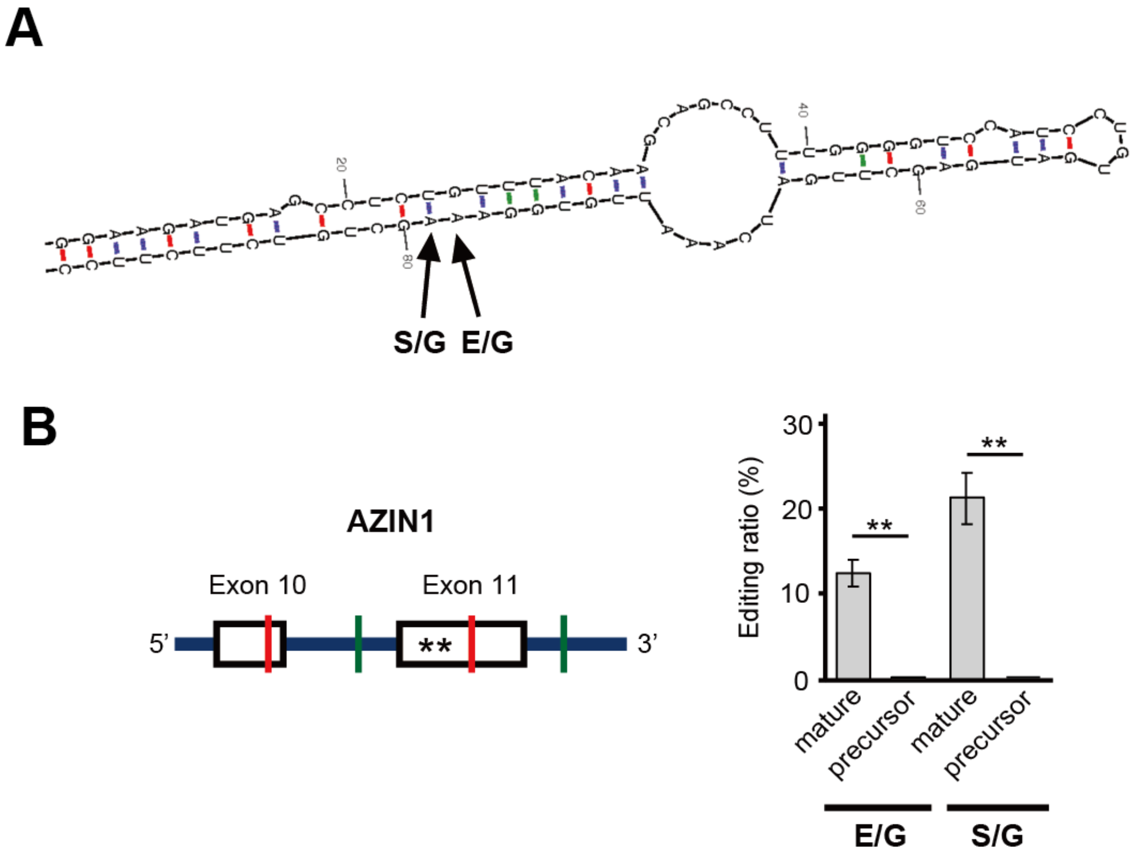


**SUPPLEMENTAL FIGURE S1. Estimated dsRNA structure required for editing at AZIN1 E/G and S/G sites.** *(A)* The secondary structure within the exon containing the AZIN1 E/G and S/G sites is estimated using Mfold. The number indicates the position from the 5' end of this exon. dsRNA, double-stranded RNA; E/G, glutamic acid/glycine; S/G, serine/glycine. *(B)* Editing ratios for the AZIN1 E/G and S/G sites in mature messenger RNA (mRNA) and precursor mRNA in the spleen of wild-type mice are shown. The locations of the two editing sites (asterisks) and primers for mature mRNA (in red) and for precursor mRNA (in green) are indicated on the left panel. Editing ratios are displayed as the mean ± SEM (n=3 mice; Student's *t*-test, $^{**}p < 0.01$).

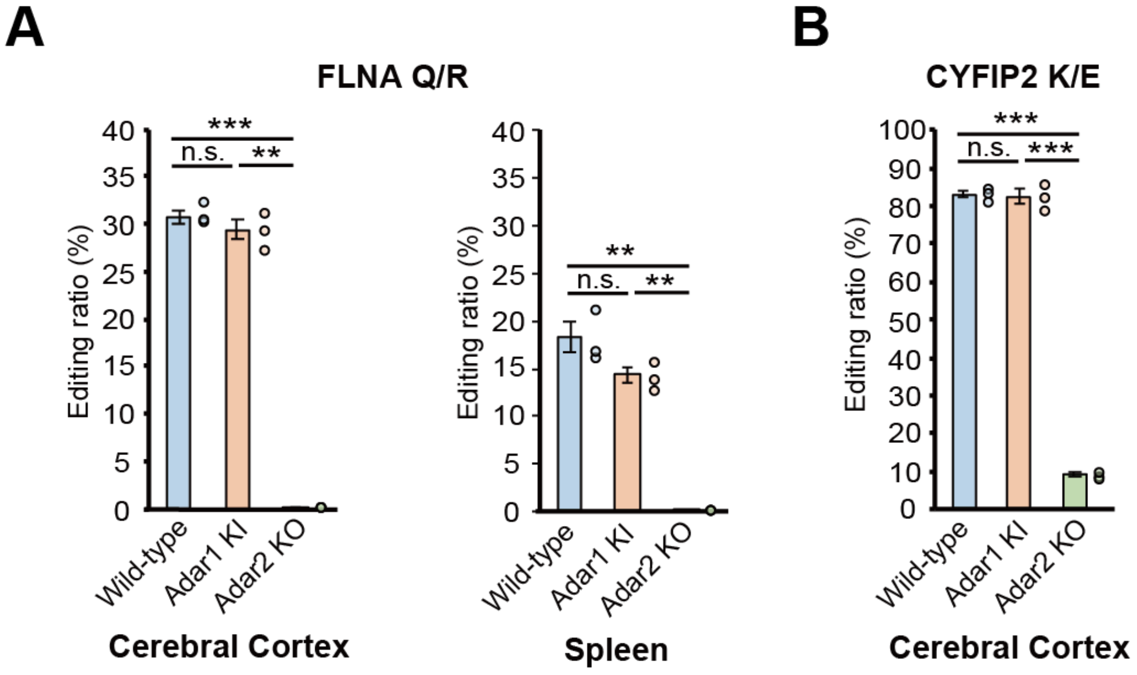


**SUPPLEMENTAL FIGURE S2. Retention of RNA editing at known ADAR2 sites in Adar1 KI and Adar2 KO mice.** (*A*-*B*) Editing ratios for the FLNA Q/R site (**A**) and CYFIP2 K/E site (*B*) in the indicated tissues isolated from wild- type (WT), *Adar1*$^{E861A/E861A}$*Ifih*$^{-/-}$ mice (Adar1 KI) and *Adar2*$^{-/-}$*Gria2*$^{R/R}$ (Adar2 KO) mice are shown. Editing ratios are displayed as the mean ± SEM (n=3 mice for each group; Student's *t*-test, $**p < 0.01$, $***p < 0.001$, n.s., not significant). The editing ratio of each mouse is also displayed as a circle on the right side.

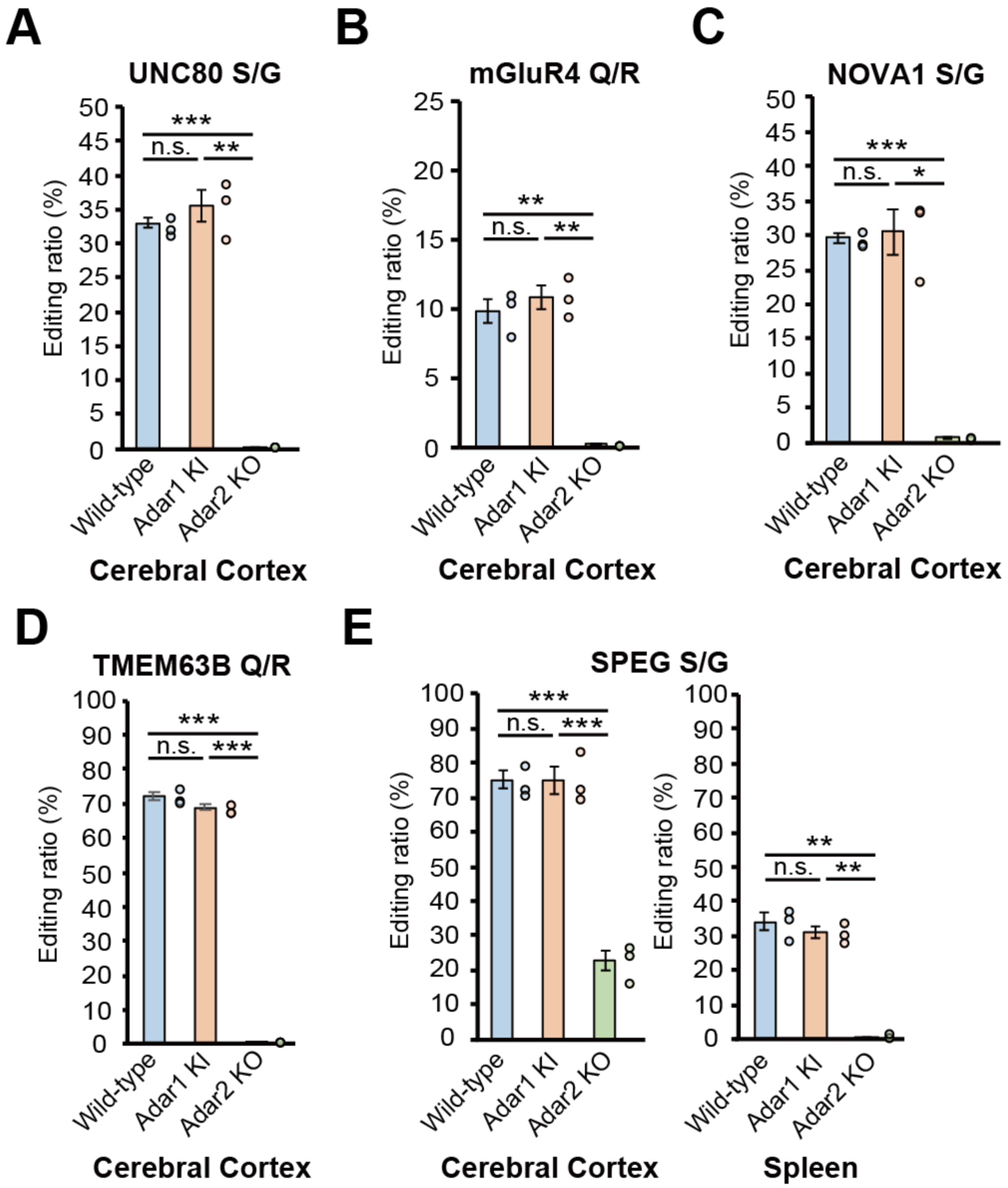


**SUPPLEMENTAL FIGURE S3. Novel ADAR2 sites.** (*A*–*E*) Editing ratios for the UNC80 S/G site (*A*), mGluR4 Q/R site (*B*), NOVA1 S/G site (*C*), TMEM63B Q/R site (*D*) and SPEG S/G site (*E*) in the indicated tissues isolated from wild-type WT, *Adar1*$^{E861A/E861A}$*Ifih*$^{-/-}$ mice (Adar1 KI) and *Adar2*$^{-/-}$*Gria2*$^{R/R}$ (Adar2 KO) mice are shown. Editing ratios are displayed as the mean ± SEM (n=3 mice for each group; Student's *t*-test, *$p < 0.05$, **$p < 0.01$, ***$p < 0.001$, n.s., not significant). The editing ratio of each mouse is also displayed as a circle on the right side. S/G, serine/glycine; Q/R, glutamine/arginine

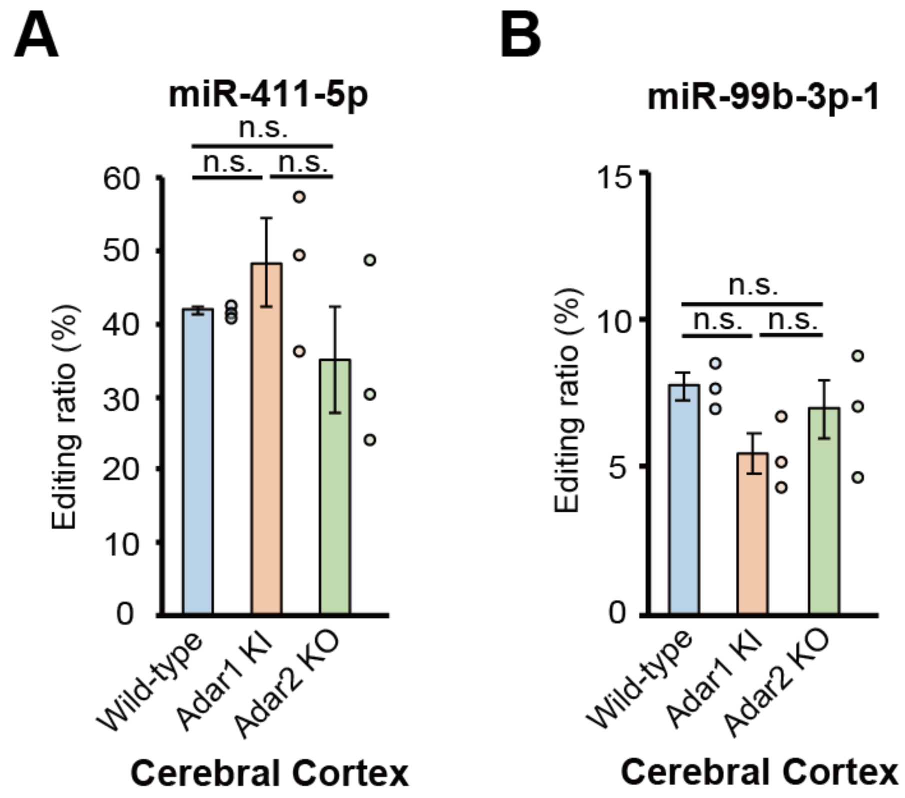


**SUPPLEMENTAL FIGURE S4. Comparable retention of RNA editing in two miRNAs from Adar1 KI and Adar2 KO mice.** (*A-B*) Editing ratios at the +5 positon of miR-411-5p (*A*) and -1 position of miR-99b-3p (*B*) in the indicated tissues isolated from wild-type (WT), *Adar1*$^{E861A/E861A}$*Ifih*$^{-/-}$ mice (Adar1 KI) and *Adar2*$^{-/-}$*Gria2*$^{R/R}$ (Adar2 KO) mice are shown. The 5' end of the mature micro (mi)RNA sequence is defined as a +1 position (See Supplemental Table S1). Editing ratios are displayed as the mean ± SEM (n=3 mice for each group; Student's *t*-test, n.s., not significant). The editing ratio of each mouse is also displayed as a circle on the right side.
